# Psi promotes *Drosophila* wing growth through transcriptional repression of key developmental networks

**DOI:** 10.1101/2020.05.13.094664

**Authors:** Olga Zaytseva, Naomi C. Mitchell, Caroline Delandre, Zuqin Nie, Maurits Evers, Janis K. Werner, John T. Lis, Ross D. Hannan, David L. Levens, Owen J. Marshall, Leonie M. Quinn

## Abstract

Psi, the sole FUSE Binding Protein (FUBP) family single stranded DNA/RNA binding protein in *Drosophila*, is essential for proper cell and tissue growth, however its mechanism of function remains unclear. Here we use Targeted DamID combined with RNA-sequencing to generate the first genome-wide binding and expression profiles for Psi. Surprisingly, we demonstrate Psi drives growth in the *Drosophila* wing through transcriptional repression of key developmental pathways (e.g. Wnt, Notch and TGFβ). Thus, Psi patterns tissue growth by directly repressing transcription of developmental growth suppressors. Analysis of direct Psi targets identified novel growth inhibitors, including *Tolkin* (Zinc metallopeptidase implicated in TGFβ signalling), *Ephexin* (Rho-GEF) and *emp* (CD36 scavenger receptor-related protein). Their depletion not only suppressed impaired growth associated with Psi knockdown, but alone was sufficient to drive wing overgrowth. Thus, Psi drives wing growth twofold, through direct activation of *Myc* and through transcriptional repression of growth inhibitors comprising core developmental pathways.

## Introduction

The human Far Upstream Binding Protein 1 (FUBP1) was isolated over a quarter of a century ago through its capacity to bind the active *MYC* promoter, remodel single stranded DNA architecture associated with RNA polymerase II activity and enable maximal activation of *MYC* transcription (Avigan et al., 1990; Duncan et al., 1994). However, the broader significance of FUBP family proteins in genome-wide transcriptional control and implications for animal development has remained unclear. *In vitro* human cell culture studies suggest FUBP1 drives *MYC* transcription in response to growth stimuli (He et al., 2000; Liu et al., 2006) and Myc expression is dysregulated in *Fubp1* knockout mouse embryonic fibroblasts (Zhou et al., 2016). The MYC oncoprotein is a potent driver of growth and cell cycle progression during development and increased MYC abundance is implicated in most human cancers (Dang, 2012; Gabay et al., 2014; Levens, 2010). Thus, understanding FUBP1-dependent mechanisms of *MYC* transcriptional control has provided insight into dysregulation of MYC in disease, particularly cancer, where elevated FUBP1 potentially leads to *MYC* overexpression and tumour progression (Debaize and Troadec, 2018). Although FUBP1 has been implicated in transcriptional control of a handful of other cell cycle control and survival genes (Debaize et al., 2018; Rabenhorst et al., 2009), whether FUBP family proteins function more widely to control transcription during animal development is currently unclear.

In *Drosophila*, the three mammalian FUBP proteins are represented by one ortholog, Psi, which also interacts with transcriptional machinery including the Mediator complex, and is required for activation of *Myc* expression, as well as cell and tissue growth during wing imaginal disc development (Guo et al., 2016). Nevertheless, the observation that *Psi* knockdown impairs cell and tissue growth in the wing more strongly than *Myc* knockdown, despite only a modest reduction in Myc abundance, suggested the downregulation of *Myc* transcription associated with Psi knockdown does not fully account for growth impairment (Guo et al., 2016). We therefore sought to determine whether Psi controls additional transcriptional targets in the wing, through intersection of Psi’s genome-wide binding profile with the Psi knockdown transcriptional signature. In addition, we aimed to determine whether altered gene expression associated with Psi knockdown might occur indirectly via Myc. To this end, we additionally identified direct targets of Myc that are differentially expressed following Myc knockdown for comparison.

Genome wide binding of Psi and Myc in the wing epithelium was determined using Targeted DamID (TaDa) (Marshall et al., 2016; Southall et al., 2013) and intersected with RNA-sequencing to detect differentially expressed genes associated with Psi or Myc depletion. Genome-wide binding profiles from TaDa revealed that, in addition to Myc, Psi directly bound genes with significant roles in development and morphogenesis. On the other hand, Myc-bound genes were uniquely enriched for roles in cell cycle and ribosome biogenesis, consistent with previously reported Myc signatures (Grewal et al., 2005; Orian et al., 2005). RNA sequencing revealed that equivalent numbers of direct Psi-targets were up- and downregulated following Psi depletion, implying that Psi not only behaves as a transcriptional activator but can also function as a repressor. Moreover, targets repressed included genes not previously implicated in growth control, revealing *tolkin* (Zinc metallopeptidase), *Ephexin* (Rho-GEF) and *emp* (CD36 scavenger receptor-related epithelial membrane protein) as novel tumour suppressor proteins. Thus, Psi modulates growth by upregulating *Myc* in concert with transcriptional repression of growth inhibitory genes comprising key developmental signalling pathways.

## Results

### Psi associates with transcriptionally active regions of chromatin

In order to identify direct Psi binding targets, we used Targeted DamID (TaDa), which was developed in *Drosophila* to exploit the GAL4/*UAS* system to determine genome-wide binding of proteins in distinct cell populations (Marshall et al., 2016; Southall et al., 2013). To identify sites of Psi enrichment specifically in larval wing discs we used the *scalloped(sd)*-GAL4 driver to express the Psi-Dam methylase. Importantly, Psi enrichment was detected on *Myc* (**Figure 1A**), thus, confirming the capacity of DamID to detect Psi’s prototypical transcriptional target. Interestingly, Psi was not only detected in proximity to the *Myc* transcription start site (TSS), consistent with roles in initiation, but was also found throughout the gene body in line with transcription elongation functions downstream of Pre-Initiation Complex (PIC) assembly. Consistent with this observation, we previously demonstrated Psi is required for enrichment of phosphorylated initiating (Ser5 CTD) and elongating (Ser2 CTD) Pol II on *Myc* (Guo et al., 2016).

**Figure 1.**
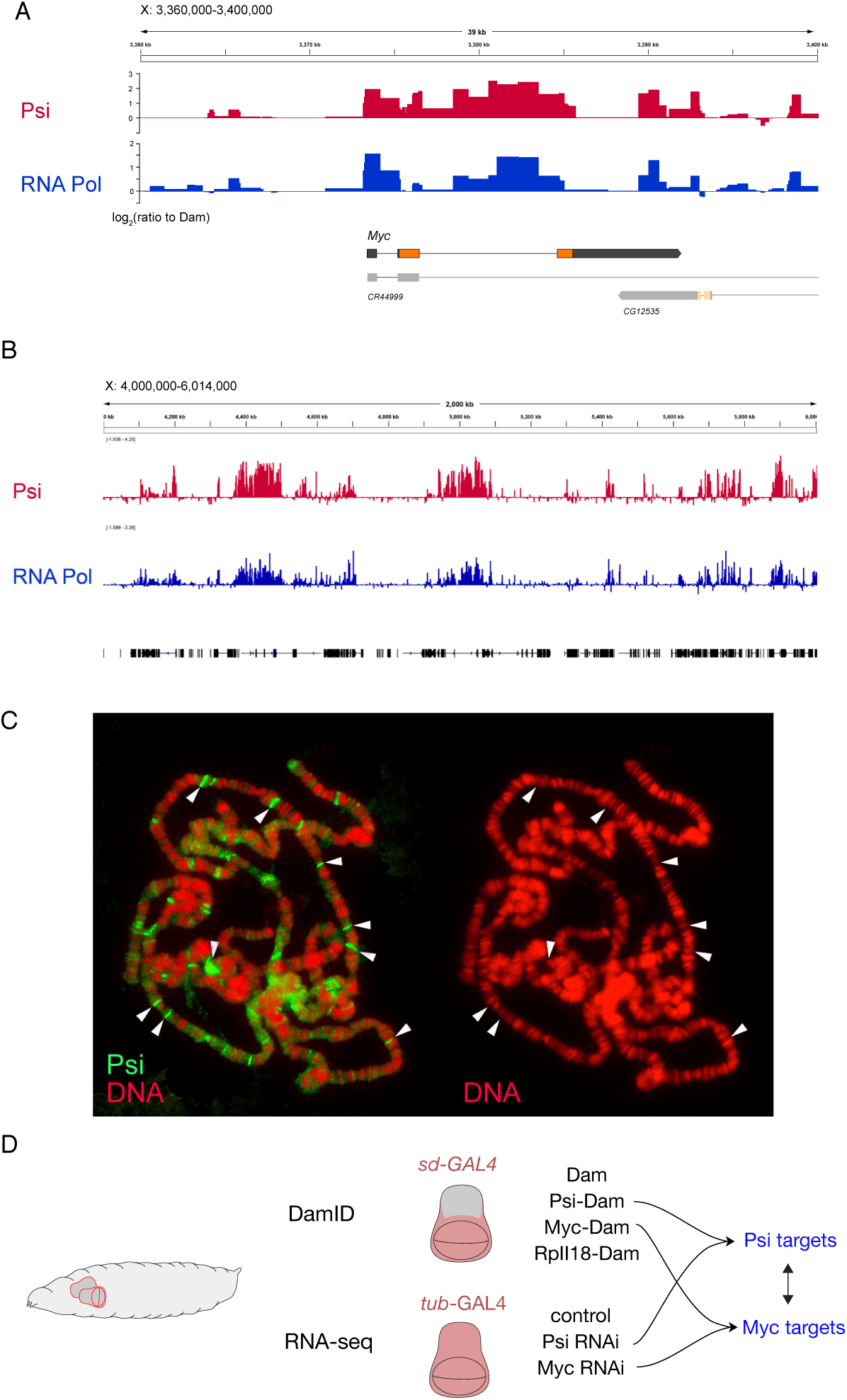
Psi binds multiple genomic regions, including Myc. (A) Psi and RNA Pol II binding profiles across the *Myc* gene in larval wing discs (*sd*-GAL4 driver used for targeted Psi-DamID and RNA Pol-DamID), shown as log_2_ of the ratio to Dam-alone control. Red: Psi binding profile; Blue: RNA polymerase binding profile. (B) Psi and RNA Pol II binding profiles across a 2 Mb region of the X chromosome. (C) Salivary gland polytene chromosomes stained with anti-Psi antibody (green). White arrows indicate regions weakly stained with DAPI (red) indicative of open chromatin. (D) Strategy to identify direct genome-wide Psi targets in wing discs. RNA-seq following Psi knockdown was used to identify differentially expressed genes, and DamID for identification of direct Psi targets. DamID using RpII18 was used to monitor transcriptional state genome-wide.

In addition to *Myc*, enrichment for Psi was detected more broadly across the genome, particularly in regions of active transcription identified based on co-enrichment with RNA Pol II (**Figure 1B**). To verify whether Psi associates with multiple genomic targets, we additionally stained polytene chromosome spreads with anti-Psi antibody. Polytene chromosomes in the *Drosophila* salivary gland arise from numerous rounds of cell endoreplication in the absence of mitosis (Edgar and Orr-Weaver, 2001). The adjacent arrangement of multiple copies of chromosomes results in DNA banding patterns, which can provide an indication of whether a protein of interest binds the genome more broadly (Johansen et al., 2009; Urata et al., 1995). Multiple bands of Psi binding were observed on polytene chromosome at sites correlating with regions weakly stained with DAPI (**Figure 1C**), indicative of open chromatin and active transcription (Lis, 2007; Pelling, 1972). Thus, in addition to *Myc*, Psi binds many direct targets at euchromatic regions.

In order to identify differentially expressed Psi targets, RNA sequencing was performed following depletion of Psi in larval wing imaginal discs, enabling intersection of binding and expression data sets (**Figure 1D**). Mammalian MYC behaves as a transcriptional amplifier, with potential to modulate cell-specific transcriptional signatures (Lin et al., 2012; Nie et al., 2012). Therefore, through regulation of Myc, Psi has the potential to indirectly regulate broad sets of target genes controlling cell and tissue growth. To determine whether Psi targets are also directly regulated by Myc, and vice versa, we concurrently identified genes differentially expressed and bound by Myc in the wing disc, using RNA-seq following *Myc* knockdown combined with Myc-DamID, respectively (**Figure 1D**). Through comparison of genome-wide Psi- and Myc- binding targets and corresponding expression profiles, we sought to determine interdependency and potential cooperation between Psi and Myc in growth regulation during wing development. Using TaDa, RNA Polymerase binding was used to map transcriptionally active genes globally (marked by the Dam-fused subunit RpII18, common to all three RNA polymerases (Filion et al., 2010)), thus, providing an indication of the transcriptional activity for Psi and Myc targets.

Prior to bioinformatic analysis of the DamID data sets, sample quality and consistency between the 3 replicates was confirmed for each condition by comparison of DamID binding profiles using pairwise Spearman correlation (**Supplemental Figure 1**). The correlation coefficient was above 0.8 for Myc and Psi, indicating sufficiently low sample-sample variability. This analysis also revealed a high level of correlation between the binding profiles for Psi and RNA Pol (ranging from 0.53 to 0.8), further suggesting Psi broadly interacts with transcriptionally active regions of the genome, as indicated by the enrichment on the *Myc* transcribed region (**Figure 1A**) and localisation of Psi to euchromatic regions of polytene chromosomes (**Figure 1C**). For further genome-wide binding analysis, the average enrichment of replicate samples was calculated at each GATC-flanked genomic fragment, to generate a representative profile.

### Genome wide Psi and Myc enrichment overlaps Pol binding

In order to visualise binding of Psi, Myc and RNA Pol binding genome-wide we compared enrichment across all gene bodies. Heatmap clustering based on Pol DamID signal into 3 groups using k-means, to identify genes with similar transcriptional activity, revealed overlap between patterns of enrichment for Pol, Psi and Myc within the 3 major gene clusters (**Figure 2A**). Cluster 1 included genes with high levels of DamID signal throughout the body of the gene and ontology analysis identified enrichment for genes implicated in ribosomal assembly and translation (**Supplemental Figure 2A**), processes of high demand in wing discs undergoing developmental growth. Cluster 2 genes were more strongly bound near transcription start sites, relative to the body of the gene (**Figure 2A**) indicative of transcriptional pausing (Adelman and Lis, 2012) and spanned a variety of functional classes including cell cycle, development and cell signalling (**Supplemental Figure 2B**). Cluster 3 genes were lowly bound by Pol, Psi and Myc (**Figure 2A**), and were enriched for neurosensory perception and mating, processes expected to be transcriptionally repressed in wing discs (**Supplemental Figure 2C**).

**Figure 2.**
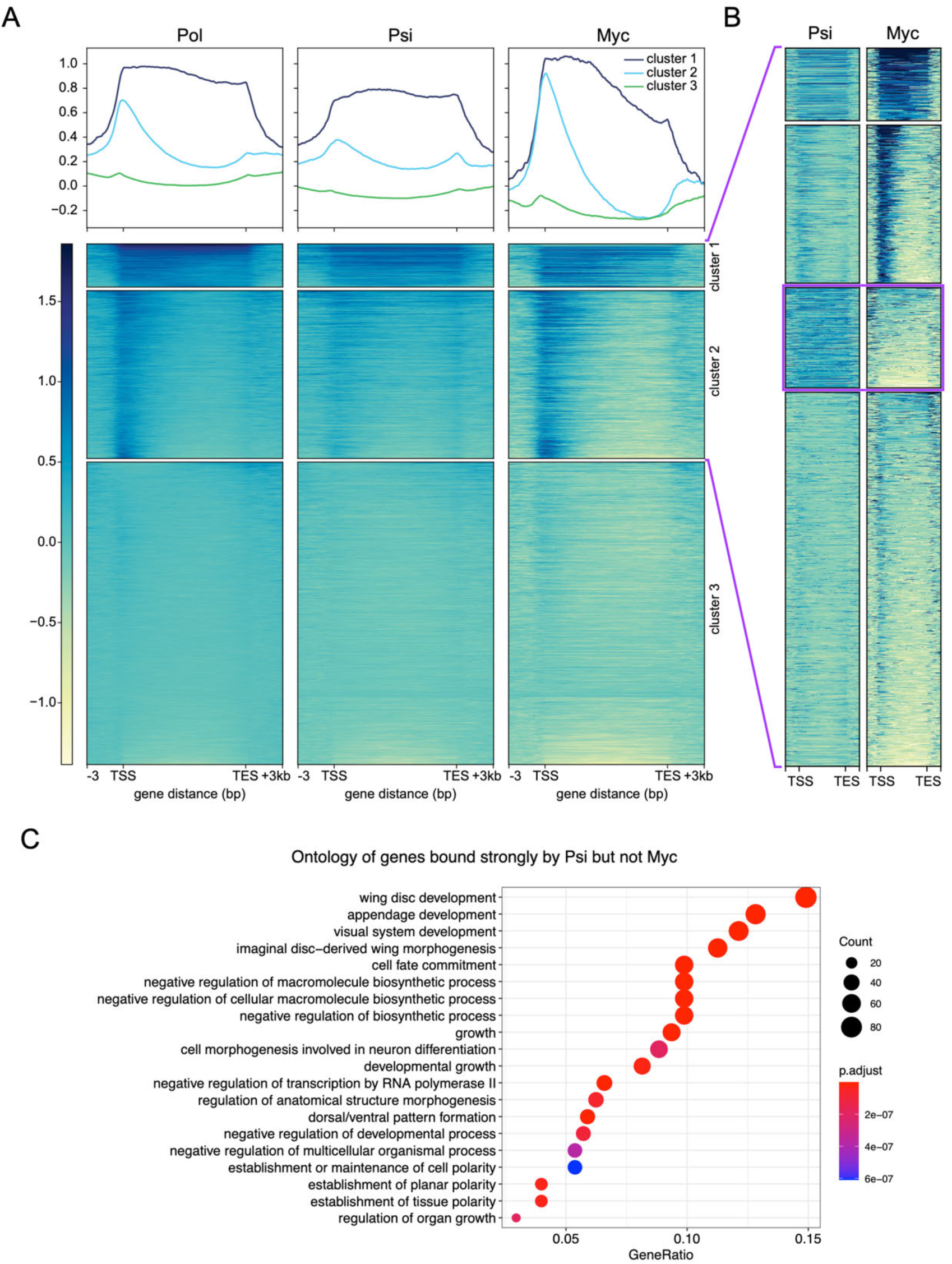
DamID binding profiles of Pol, Myc and Psi. (A) Average Pol, Psi and Myc binding across genic regions and heatmap of DamID signal, clustered by k-means into 3 clusters using Pol signal. (B). K-means clustering of Psi and Myc signal across transcriptionally active genes. (C) Ontology analysis of the gene cluster associated with binding of Psi, but not Myc, highlighted in (B).

In order to compare Psi and Myc binding profiles, the genes exhibiting low levels of DamID signal (cluster 3 in **Figure 2A**) were excluded, and the transcriptionally active genes grouped based on Psi and Myc profiles. Interestingly, clustering revealed weak Myc signal for genes sets significantly bound by Psi (**Figure 2B**). Ontology analysis of the 1023 genes in the Psi strong/Myc weak cluster identified enrichment for developmental and morphogenesis genes (**Figure 2C**), indicating Psi associates with genes involved in developmental processes independently of Myc.

### Comparison of Psi and Myc target gene functions

Peak calling on the Psi and Myc DamID profiles revealed 1449 total loci bound by Psi and 3110 loci bound by Myc, of which 908 were bound by both Psi and Myc (**Supplemental Figure 3A**). We therefore performed ontology analysis for targets bound by Psi or Myc alone and genes bound by both Psi and Myc. As expected, given the capacity of Myc to drive proliferative cell growth (Grewal, Li, Orian, Eisenman, & Edgar, 2005; Johnston, Prober, Edgar, Eisenman, & Gallant, 1999; Wu & Johnston, 2010), ribosome biogenesis and cell cycle genes were enriched among genes bound by Myc (**Supplemental Figure 3B**). These processes were not, however, enriched in the Psi binding dataset, which comprised genes implicated in cell fate, development, morphogenesis and signalling (**Supplemental Figure 3B**). Enrichment for growth, development, morphogenesis and signalling pathway regulatory genes was also observed for genes bound by both Myc and Psi (**Supplemental Figure 3B**). Together these data suggest Myc regulates transcriptional networks driving accumulation of biomass independently of Psi. Conversely, the growth impairment observed in wings following *Psi* knockdown (Guo et al., 2016) occurs via both Myc-dependent and Myc-independent targets.

### Genome-wide analysis of Psi and Myc expression signatures using RNA-seq

To identify genes with significantly altered expression following *Psi* knockdown, we performed RNA sequencing of larval wing imaginal discs. Principal component analysis revealed separation between genotypes and consistency between replicates (**Supplemental Figure 4**). Analysis of differential gene expression compared with control wings revealed that, at a False Discovery Rate (FDR) cut-off of 1%, 2347 genes were differentially expressed in *Myc* RNAi wings (**Figure 3A**) and 882 in *Psi* RNAi wings (**Figure 3B**). Consistent with previous observations, *Myc* mRNA levels were significantly reduced after knockdown of *Psi* (**Figure 3B**, log_2_FC=-0.369, adjusted p-value=0.0008), indicating capacity of RNA-seq to detect differential expression of this verified Psi target (Guo et al., 2016). Within the *Psi* knockdown dataset, 429 genes were downregulated and 453 upregulated, suggesting Psi has capacity to both up- and down-regulate expression of targets (**Figure 3B**). The majority of genes significantly altered following *Psi* knockdown displayed less than 2-fold changes, indicating most genes require Psi for fine-tuning, rather than acute activation or repression of expression. This observation is in accordance with the predicted function of Psi’s mammalian counterpart, FUBP1, as a “cruise control” in the modulation of active *MYC* transcription (Levens, 2013; Liu and Levens, 2006; Zaytseva and Quinn, 2017).

**Figure 3.**
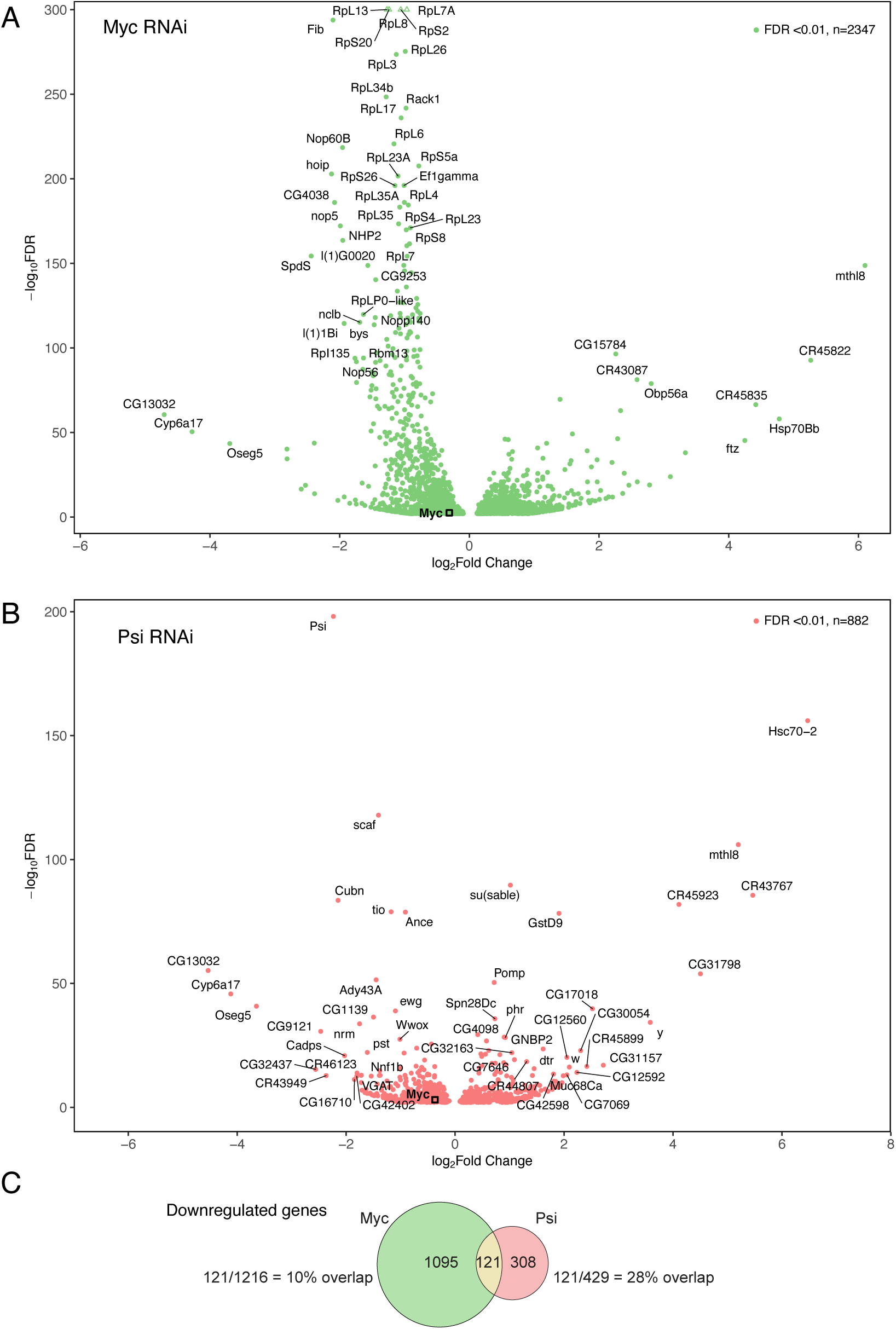
Significantly altered genes following Myc or Psi knockdown in wing discs. Genes with statistically significant altered expression following (A) Myc knockdown and (B) Psi knockdown at FDR<0.01 are shown. Top 50 genes with greatest fold change and smallest p-value are labelled. *Myc* is highlighted with a black square in each plot. (C) Intersection of gene sets downregulated after either Myc or Psi knockdown.

Mammalian MYC behaves as a global amplifier of gene expression, with increased levels of MYC leading to an overall increase in expression of transcriptionally active genes (Lin et al., 2012; Nie et al., 2012). Given the high degree of functional conservation between *Drosophila* and mammalian MYC (Schreiber-Agus et al., 1997), we predicted that *Drosophila* Myc would also function as a transcriptional amplifier. In support of this idea, microarray-based approaches investigating short-term *Drosophila* Myc overexpression, induced by heat shock in third instar larval tissues, identified broad increases in expression of endogenous target genes (Orian et al., 2003). We therefore first analysed genes with decreased expression in Myc depleted wings, identifying 1216 genes significantly downregulated after *Myc* knockdown (**Figure 3A**). In accordance with ontology classes determined by DamID-seq (**Supplemental Figure 3B**), analysis of downregulated genes in Myc-depleted wing discs identified ontologies largely associated with cell and tissue growth, including ribosome biogenesis and cell cycle control (**Supplemental Figure 5**). Thus, as expected, given the cell growth signature associated with Myc overexpression (Grewal et al., 2005), we demonstrate Myc depletion impairs expression of genes required for proliferative cell growth, which are highly active in growing tissues, such as the developing wing.

Intersection of gene sets downregulated in *Myc* or *Psi* knockdown wing discs revealed 121 overlapping targets (**Figure 3C**). The relatively small overlap between differentially expressed genes (10% of Myc genes, 28% of Psi genes) suggests more than 2/3rds of candidate Psi targets are regulated by Psi independently of Myc (**Figure 3C**). Interestingly, the overlap between the Psi and Myc transcriptome, identified by RNA-seq alone (28%, **Figure 3C**), was smaller than the 63% percent of genes co-bound by Psi and Myc based on DamID analysis (**Supplemental Figure 3A**). Binding events occurring without altered expression may be explained by the temporal differences between the analyses, where binding using Dam methylation was detected over a 24-hour window, while changes in gene expression were detected 3 days after inducing knockdown. Thus, over the 3-day window, RNA-seq will detect indirect changes to rates of transcription and/or stabilisation of target mRNA species enacted through multiple feedback mechanisms.

### Psi regulates gene expression levels independently of splicing

In addition to binding ssDNA, mammalian FUBP-family proteins also bind RNA via their KH domains to regulate RNA processing (Gherzi et al., 2004; Miro et al., 2015). Psi also binds RNA via the KH motifs to control RNA splicing (Brooks et al., 2015; Labourier, Blanchette, Feiger, Adams, & Rio, 2002; Q. Wang et al., 2016). Therefore, to determine whether splicing functions might augment Psi’s transcriptional roles in wing discs, we performed analysis of differential splicing following *Psi* knockdown with rMATS (Shen et al., 2014), which identifies mis-spliced events and additionally has the capacity to discover unannotated splice sites. rMATS detected 1349 events at 582 genes with differential splicing. Classification into splicing event types by rMATS identified exon skipping (53%) and mutual exon exclusivity (26%) as the most common alterations in Psi knockdown (**Supplemental Figure 6A**).

Ontology analysis of differentially spliced genes following *Psi* knockdown revealed enrichment for developmental processes (**Supplemental Figure 6B**). Thus, Psi may regulate development both through altered expression of target genes and via modulation of splicing. Intersection of expression and splicing data sets revealed 111 genes both differentially expressed and alternatively spliced (**Supplemental Figure 6C**). The relatively small overlap (13% of alternatively spliced genes and 19% of differentially expressed genes) indicates most transcriptional and splicing alterations occur independently. Thus, defective coupling of transcription and splicing, where impaired transcription indirectly alters splicing patterns (Bentley, 2014), is unlikely to explain these observations. Moreover, intersection of the alternatively spliced genes with the Psi binding profile indicated that the majority (71%) of these events did not require association of Psi with the transcribed region of the gene (**Supplemental Figure 6D)**.

Importantly, *Myc* was not differentially spliced following *Psi* knockdown based on the rMATS analysis, as *Myc* downregulation occurred without a relative change in the proportion of reads overlapping the introns (**Supplemental Figure 6E**). Thus, Psi predominantly functions to regulate *Myc* at the level of transcription. Together, the direct interaction between Psi and the *Myc* gene (**Figure 1A**), the requirement for Psi in RNA Pol II loading on *Myc* and maintenance of *Myc* mRNA levels (Guo et al., 2016), strongly suggests that Psi is required for direct regulation of *Myc* transcription rather than RNA processing of *Myc*.

### Direct targets of Psi function in development

Next, we intersected direct Psi targets with genes significantly altered following *Psi* knockdown. Only 153 genes were shared between the two gene sets (**Figure 4A**,**B**). Thus, 73% of genes with altered expression after Psi knockdown are likely regulated indirectly, via a downstream transcriptional regulator and/or post-transcriptionally (**Figure 4A**). Furthermore, significant differential gene expression was not observed for the majority (89%) of genes bound by Psi. Therefore, the combination of transcriptome and binding profiling was essential for identification of Psi’s direct and differentially expressed targets. Ontology analysis of these direct and differentially expressed targets revealed that Psi modulates genes implicated in growth, development and morphogenesis (**Figure 4C**), including development of the wing (**Figure 4D**)

**Figure 4.**
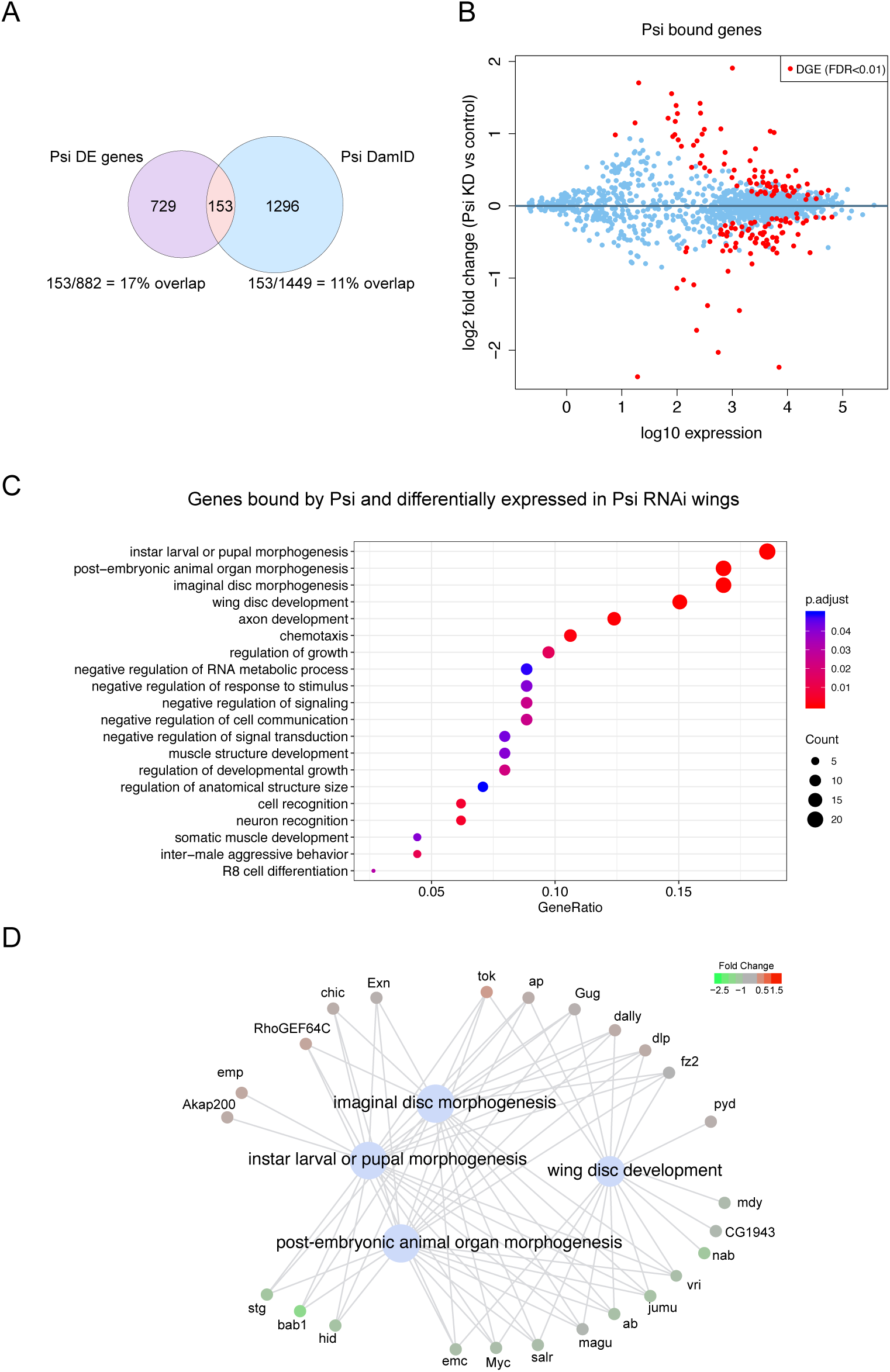
Psi binds and regulates developmental genes. (A) Intersection of differentially expressed genes after Psi knockdown with genes bound by Psi. (B) MA plot showing only genes bound by Psi (blue) while statistically significant DGE events at FDR<0.01 are shown in red. (C) Ontology of mutually inclusive genes from the intersection in (A). (D) Genes regulated by Psi with known roles in wing morphogenesis.

### Knockdown of genes repressed by Psi restores Psi-dependent growth

We next sought to identify Psi targets, determined by RNAseq and DamID, required for Psi-dependent growth. We have previously shown *Myc* transcriptional activation requires Psi (Guo et al., 2016), however, many direct Psi targets were upregulated following *Psi* knockdown suggesting a novel role for Psi as a transcriptional repressor. We hypothesized depletion of genes bound by Psi and upregulated in *Psi* knockdown wings would rescue the impaired growth phenotype. We therefore used RNAi transgenes to co-deplete direct Psi-repressed targets *Akap200, emp, Exn, chic, dally, dlp, fz2* and *tok* (**Figure 4D**), in Psi depleted wings (knockdown validated by qPCR **Figure 5A and Supplemental Figure 7A**). Impaired wing growth associated with *Psi* depletion was not altered by co-knockdown of *Akap200, chic* nor *dally* (**Supplemental Figure 7B**). However, knockdown of *emp, Exn* and *tok* not only suppressed the Psi small wing phenotype (**Figure 5B**) but their depletion alone was sufficient to drive overgrowth (**Figure 5B**), providing evidence emp, Exn and tok normally function as negative regulators of wing growth. Thus, increased levels of *emp, Exn* and *tok* expression associated with Psi knockdown, detected by RNA-seq, likely contribute to impaired growth in the Psi-depleted wing.

In contrast to the other genes tested, *fz2* knockdown alone did not alter wing size (**Figure 5B**), rather, the impaired growth caused by Psi depletion was dependent on fz2, as co-knockdown suppressed the *Psi* RNAi small wing phenotype (**Figure 5B**). fz2 (frizzled2) is a transmembrane receptor for the wg ligand (Wnt homologue in *Drosophila*), which binds wg to control cell cycle exit and differentiation of cells comprising the larval wing margin (Johnston & Edgar, 1998; Zhang & Carthew, 1998). More recent reports revealed fz2 is also expressed in the periphery of the larval wing pouch and required for long-range Wg signalling and cell survival (Chaudhary et al., 2019). The observation that impaired growth associated with Psi knockdown requires fz2, while fz2 knockdown alone does not alter wing growth, suggests networks modulated by Psi require Wg pathway activity. Thus, Psi controls wing growth by functioning as a transcriptional repressor of several developmental determinants. Moreover, Wg signalling is required for Psi-dependent growth.

**Figure 5.**
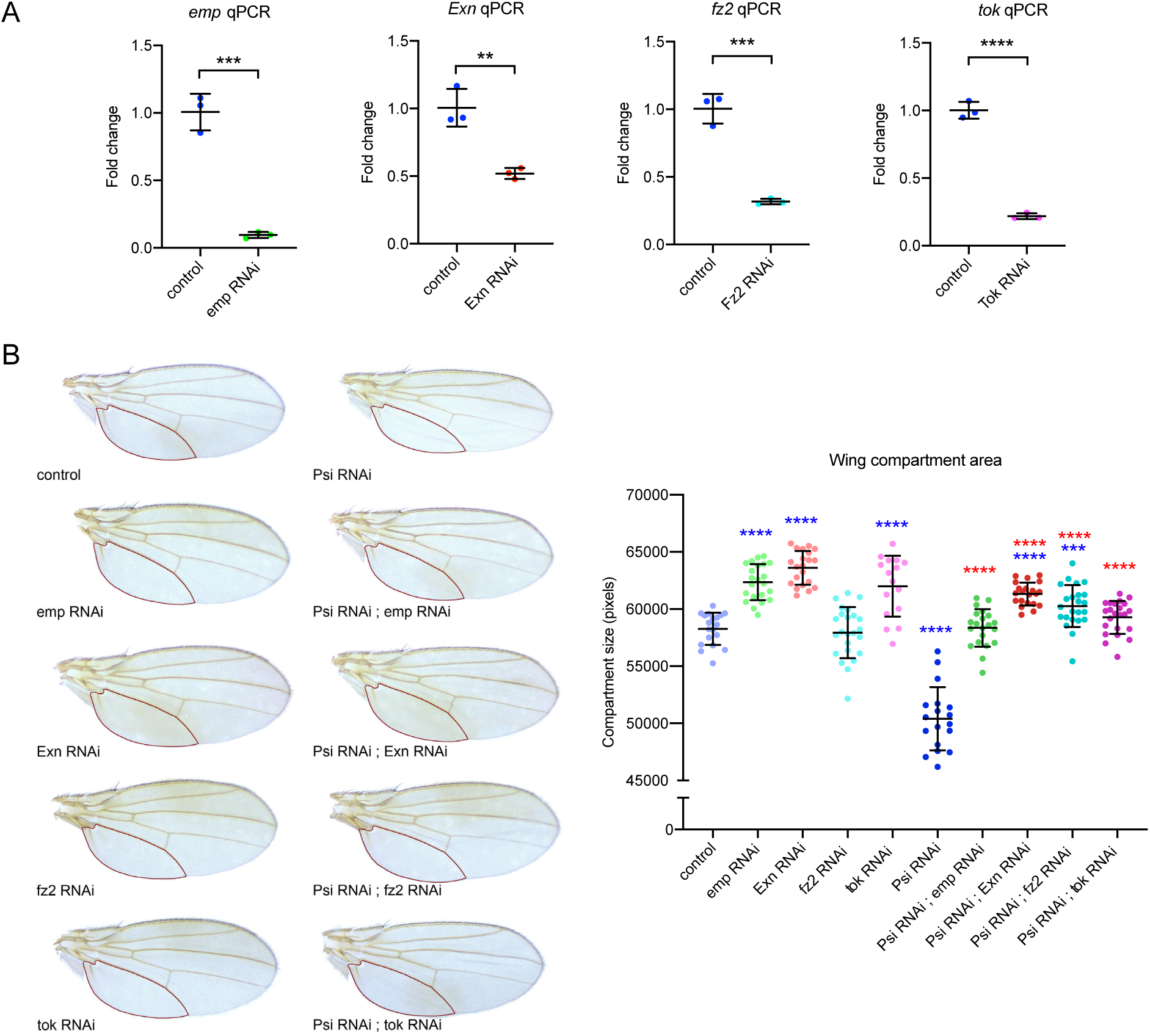
Targets negatively regulated by Psi are required for Psi-dependent growth. (A) qPCR of third instar larval wing discs 2 days after induction of RNAi transgenes (using tsGAL80; *tub*-GAL4) for Psi targets *Emp* (p=0.0003), *exn* (p=0.0044), *fz2* (p=0.0005) and *tok* (p<0.0001). (B) Adult wings with *ser*-GAL4 driven knockdown of Psi targets required for wing growth. Quantification of the posterior compartment of the adult wing defined by the L5 vein, marked with the red outline. P-values were corrected for multiple testing using the Benjamini-Hochberg FDR method (blue stars indicate comparison to control; red stars indicate comparison to *Psi* RNAi). **** indicates p_adj_<0.0001, *** p_adj_=0.0005.

## Discussion

Here we report the genome wide binding signature for Psi, which displays strong overlap with active chromatin. Ontology analysis of direct and differentially expressed Psi targets, suggest Psi functions to modulate transcription of proliferative growth during development. Psi also functions as a splicing regulator (Labourier et al., 2002; Wang et al., 2016), however, Psi’s transcriptional and splicing functions appear largely independent in the wing with limited overlap between differentially expressed and spliced genes. Thus, although transcription and splicing are often tightly coupled, such that defective transcription can indirectly alter splicing (Bentley, 2014), this is unlikely in the case of *Psi* knockdown. Importantly, splicing changes were not observed for direct targets implicated in Psi-dependent growth.

Together our data demonstrate that Psi potentiates tissue growth by directly modulating expression of key components of multiple developmental pathways. The model for Psi-mediated gene regulation is summarised in **Figure 6**. Psi binds and modulates *Myc* transcription, however, transcriptional signatures associated with *Psi* or *Myc* knockdown show limited overlap. The predominant Myc signature is ribosome biogenesis and proliferative growth, processes not enriched for Psi, suggesting growth impairment associated with Psi depletion is unlikely a direct consequence of Myc function. Rather, Psi directly binds and modifies expression of developmental growth patterning genes. The direct targets of Psi transcriptional repression, *Exn, emp* and *tok*, are essential growth inhibitors in the wing and also required for the growth impairment associated with *Psi* knockdown. Moreover, through their identification as direct Psi targets, we provide the first evidence that emp and tok function as novel growth suppressors.

**Figure 6.**
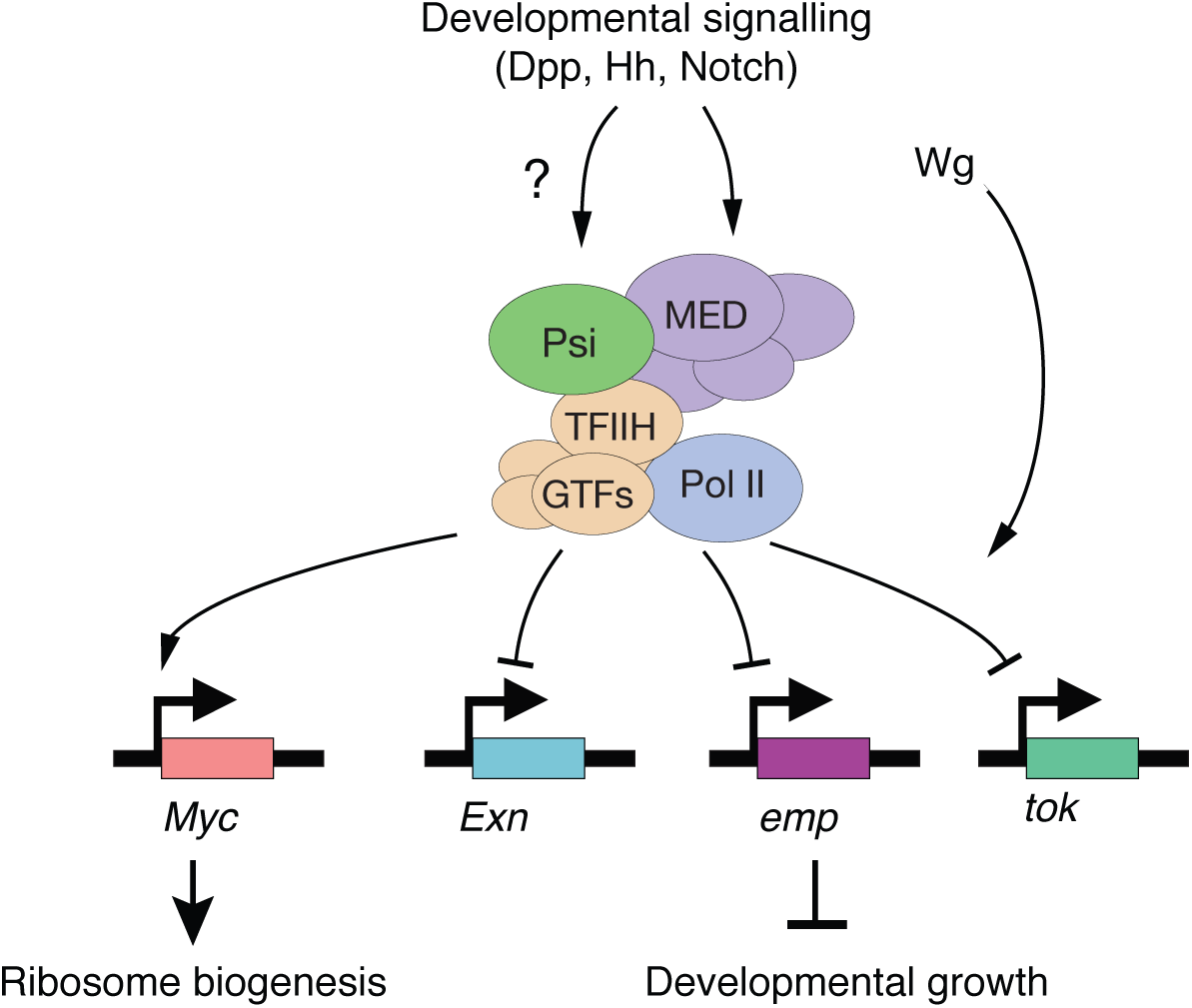
Model for regulation of wing development by Psi.

*Exn* (*Ephexin*) encodes a Rho Guanine nuclear exchange factor (RhoGEF) implicated in imaginal disc morphogenesis, including formation of leg joints (Greenberg & Hatini, 2011). In the context of the wing, genome-wide association analysis of single nucleotide polymorphisms (SNPs) associated with altered body size and wing growth identified clusters of SNPs within the *Exn* locus, while wing-specific knockdown of *Exn* via RNAi resulted in wing overgrowth (Vonesch, Lamparter, Mackay, Bergmann, & Hafen, 2016), consistent with our observation that Exn behaves as a growth inhibitor. Moreover, co-depletion of *Exn* suppressed the Psi small wing phenotype, suggesting *Exn* upregulation mediates the impaired growth associated with Psi knockdown.

emp (epithelial membrane protein) is expressed on the surface of epithelial cells in larval wing imaginal discs (Hart & Wilcox, 1993), however, emp function(s) in the wing are currently unknown. *emp* mRNA is upregulated in embryonic lipid accumulation stages during *Drosophila* oogenesis with unknown significance (Sieber & Spradling, 2015). emp possesses homology with mammalian CD36, a glycoprotein that responds to lipids, glycoproteins and lipoproteins to regulate angiogenesis and fatty acid uptake. CD36-null mice display elevated ovarian angiogenesis and increased proliferation of follicular cells (Osz et al., 2014) and human CD36 may have potential as a prognostic marker for metastatic cancer of epithelial origin (Enciu, Radu, Popescu, Hinescu, & Ceafalan, 2018; Ladanyi et al., 2018; Nath, Li, Roberts, & Chan, 2015). Here we demonstrate *emp* loss-of-function suppresses the Psi phenotype, which suggests direct repression of *emp* transcription by Psi is normally required to constrain wing growth. Further to this, we show *emp* depletion is sufficient to drive wing overgrowth, however, further studies are required to determine whether the tumour suppressor function of emp is dependent on lipogenesis or fatty acid uptake.

The BMP-1 homologue tok (tolkin) regulates Dpp signalling in early embryogenesis (Finelli, Xie, Bossie, Blackman, & Padgett, 1995) and also cleaves the secreted BMP antagonist sog, to drive cross vein formation in the pupal wing (Serpe, Ralston, Blair, & O’Connor, 2005). Although Serpe et al (2005) reported vein defects using *tok* loss-of-function mutants, loss of wing veins was not observed after *ser*-GAL4 mediated knockdown, likely due to continued production and secretion of tok from ventral compartment cells enabling vein formation. Here we provide the first evidence the tok metalloproteinase is an essential growth inhibitor in the wing, suppressing the Psi small wing phenotype and demonstrating depletion alone was sufficient to drive wing overgrowth.

Future studies investigating the mechanisms linking cellular signalling via emp, Exn, and tok pathways to proliferative growth control will be important. In particular, in contrast to Psi targets required for the Psi knockdown phenotype (i.e. *emp, Exn*, and *tok*), *fz2* depletion alone did not alter wing size. However, Psi knockdown was unable to decrease wing size in the *fz2* depleted background, suggesting cellular targets mis-regulated by Psi knockdown required for wing growth require Wg signalling.

Previous analysis of Psi’s protein interactors revealed components of the Mediator (MED) complex (Guo et al., 2016), which modulates context-dependent transcriptional networks by sensing cellular signalling programs (Allen and Taatjes, 2015; Carrera et al., 2008; Kim et al., 2006; van de Peppel et al., 2005; Zhao et al., 2013). Interestingly, in addition to MED, one of Psi’s top protein interacting partners is boca, an endoplasmic reticulum protein required for trafficking of the LDLR family receptor arrow (arr), which together with either fz or fz2 is activated by the wg ligand (Culi and Mann, 2003; Wehrli et al., 2000). Furthermore, Psi physically interacts with disheveled (dsh), a conserved Wg pathway adaptor; upon activation of fz/fz2, dsh sequesters the APC/Axin protein destruction complex, allowing stabilization of β-catenin (armadillo in *Drosophila*) and activation of Wg-transcriptional targets (reviewed in (Bejsovec, 2006)). Thus, Psi may connect key components of signalling pathways, including Wg, with the MED complex to confer transcriptional specificity for MED-activation of defined developmental targets in response to upstream signals. Given that Psi also controls expression of the Dpp pathway, which has capacity to indirectly modulate expression of downstream Wg targets (Yang, Meng, Ma, Xie, & Fang, 2013), further studies investigating interplay between these major developmental signalling pathways and Psi will be of great interest.

In summary, Psi fine tunes transcription of multiple signalling networks to coordinate growth control during wing development. Interestingly, although FUBP1-like proteins have been annotated in *C. elegans* (Davis-Smyth et al., 1996), similar proteins are not present in yeast. The FUBP-family may, therefore, have arisen to enable patterning of cell growth, proliferation and differentiation, which is essential for development of all multicellular organisms.

## Materials and Methods

### Expression constructs

*pTaDaG-Psi* and *pTaDaG-Myc* were generated by PCR amplifying the ORF inserts from DRGP plasmids FMO12803 (*Myc*) and FMO09121 (*Psi*) and cloning into the *pTaDaG* vector (Delandre, McMullen, & Marshall, 2020) cut with BglII/XhoI via NEB HiFi Assembly (NEB). PCR primers for NEB HiFi Assembly were designed using PerlPrimer (Marshall, 2004). *pTaDaG-RpII18* was generated via the insertion of a custom gBlock (IDT) containing *cMycNLS-linker-RpII18-RA* ORF into *pTaDaG* cut with BglII/XhoI via NEB HiFi Assembly. Primer and gBlock sequences provided in Supplemental Information.

### Fly stocks

Fly stocks were maintained on a standard molasses and semolina *Drosophila* medium. Genetic crosses were raised at 25°C except when performed in the background of temperature-sensitive GAL80 expression, where they were initially raised at 18°C followed by a shift to 29°C. The *Serrate-*GAL4 (BL6791), *Scalloped-*GAL4 (BL8609), *Tubulin-*GAL4 (BL5138), *Tubulin-* GAL80^ts^ (BL7019), *UAS-Exn* RNAi (BL33373), *UAS-fz2* RNAi (BL31312), *UAS-chic* RNAi (BL34523), *UAS-dally* RNAi (BL33952), *UAS-dlp* RNAi (BL34091) lines were obtained from the Bloomington *Drosophila* Stock Centre. The *UAS-Psi* RNAi (v105135), *UAS-Myc* RNAi (v2947), *UAS-emp* RNAi (v12233), *UAS-tok* RNAi (v2626), *UAS-Akap200* RNAi (v102374) were obtained from the Vienna *Drosophila* Resource Centre.

Targeted DamID lines generated for this study (*TaDaG-psi, TaDaG-myc, TaDaG-rpII18*) were generated by BestGene, Inc (CA), through phiC31-integrase-mediated insertion of the appropriate expression vectors into attP2 on chromosome 3. TaDaG-Dam flies were used as previously described (Delandre et al., 2020).

### Polytene immunostaining

The larvae were heat shocked for 20 min at 36.5°C. Polytene squashes and immunofluorescence labeling was done as previously described (Schwartz et al., 2004). The chromosomes were stained with Psi antibody (raised against full-length Psi protein in guinea pigs) at 1:20 and Hoechst 33258 for labeling of DNA. The fly line used was WTD7(87E) from David Gilmour (Wu et al., 2003).

### DamID sample preparation

Embryos from parental crosses using the *sd*-GAL4; *tub*-GAL80^ts^ driver were collected over the course of <4-hour lays at 25°C, after which the embryos were placed at the repressive temperature of 18°C for 7 days until the second larval instar stage. The larvae were then shifted to the permissive temperature of 29°C for 24h. Larval wing discs were collected into cold PBS, genomic DNA was extracted using a Zymo Quick-DNA kit (#D4069) after treatment with Proteinase K for 1-3 hours at 56°C in the presence of 50 µM EDTA. GATC methyl-specific digest using DpnI was carried out at 37°C overnight, and cleaned up using a Machery-Nagel PCR purification kit (740609.50). Samples were eluted into 30 µL H_2_O and 15 µL was used for subsequent preparation. Adaptors for PCR enrichment of methyl-digested sites were ligated for 2 hours at 16°C using T4 DNA ligase. Digest of unmethylated GATC sequences was performed with DpnII at 37°C for 2 hours, in order to decrease signal from unlabelled sites. PCR using MyTaq polymerase (Bioline BIO-21113) was performed with 3 long extension cycles followed by 17 short extension cycles as described (Vogel, Peric-Hupkes, & van Steensel, 2007). The PCR products were cleaned up again with a Machery-Nagel PCR purification kit. PCR adaptors were removed by overnight digest at 37°C with AlwI. Samples were sonicated in 100 µL volumes using a Covaris S2 sonicator at 10% DUTY, 140W peak incident, 200 cycles per burst, 80 second duration, achieving 300 bp average fragment size. Sample clean-up and library preparation was carried out using Sera-Mag Speedbeads hydrophobic carboxyl magnetic beads (GE Healthcare, 65152105050250). Following bead cleanup, sample concentrations were measured using Qubit DNA HS reagents (Thermo Fisher, Q32854) and <500 ng of DNA for each sample was used to generate the libraries. End repair was performed for 30 min at 30°C with T4DNA Polymerase, Klenow Fragment and T4 polynucleotide kinase. 3’ ends were adenylated using Klenow 3’ to 5’ exo-enzyme for 30 min at 37°C. Unique index adaptors were ligated to each sample using NEB Quick Ligase for 10 min at 30°C. The samples were cleaned up with Sera-Mag beads twice to ensure the removal of sequencing adaptor dimers. DNA fragments were enriched by PCR using NEB Next Ultra II Q5 Master Mix (NEB M0544S), before final clean up using Sera-Mag beads. Successful ligation of adaptors and the absence of adaptor concatemers were verified using an Agilent Bioanalyser, and the final concentration was measured using Qubit. The libraries were pooled to achieve an equimolar concentration of each sample based on average fragment size and concentration, with the final total concentration of 2 nM. The samples were sequenced using HiSeq2500 Illumina platform in Rapid Run mode, with 50 bp single-end reads.

### DamID analysis

The DamID dataset was analysed using a single pipeline workflow (Marshall & Brand, 2015). The damidseq_pipeline script was used to align the reads to the *Drosophila* BDGP6 genome with Bowtie2, identify GATC sites and calculate the normalised log_2_ ratio between Dam-fusion protein profile and Dam alone. Spearman sample correlation and genomic coverage clustered metaplots were generated using the deepTools package (Ramírez et al., 2016), using the output of the damidseq_pipeline bedgraph files converted into bigwig files. Average enrichment between replicates were determined by calculating average coverage at each region flanked by GATC sites. Enrichment profiles in bedgraph format were visualised using the Integrative Genome Viewer (IGV). Significant peaks were detected at 1% FDR using the find_peaks script, peaks2genes script to identify genes within 1kb of the discovered peaks and transcriptionally active genes were identified using the polii.gene.call script (Marshall & Brand, 2015).

### RNA-seq

Larval wing discs were collected after 3 days of GAL4-induced knockdown. For each sample, 3 collections of 20 larval wing discs were pooled (60 wing discs in total). RNA was extracted using the Promega ReliaPrep RNA Tissue miniprep system and eluted in 50 µL nuclease-free water and RNA integrity verified using a Bioanalyser Tapestation. Library preparation was carried out by the ACRF Biomolecular Resource Facility, John Curtin School of Medical Research, Australian National University. RNA was prepared using the standard TruSeq Illumina protocol preserving strandedness information, with Oligo-dT beads used to enrich for mRNA and exclude other RNA. Samples were sequenced using the HiSeq2500 Illumina system, with 100 bp paired-end reads.

### Differential expression analysis

RNAseq sequences were aligned to the *Drosophila melanogaster* genome Flybase release 6.10 using Tophat2. The gene counts were performed using HTSeq Python package (Anders, Pyl, & Huber, 2015). Significant differential expression was analysed using DESeq2 R package (Love, Huber, & Anders, 2014), with FDR cutoff 1% used to identify statistically significant events.

### Gene ontology analysis

Gene Ontology analysis of Entrez IDs associated with significantly altered genes was performed using the clusterProfiler R package (Yu, Wang, Han, & He, 2012). The Benjamini-Hochberg multiple testing correction method was used and adjusted p-value cutoff of 0.05 was applied. The clusterProfiler filtering function was applied to exclude parent terms, where applicable. Gene ontology networks of the clusterProfiler output were generated in Cytoscape using the EnrichmentMap plugin (Merico, Isserlin, Stueker, Emili, & Bader, 2010), and gene ontologies were manually grouped and annotated based on similarity.

### Differential splicing analysis

Analysis of differential splicing was performed using rMATS 4.0.2 (Shen et al., 2014) on BAM files aligned for differential expression. Junction reads as well as reads covering the exon of interest were used to calculate differences in exon inclusion rates. Adjusted p-value cutoff of 0.01 was applied to detect significant splicing changes. The ggsashimi package (Garrido-Martín, Palumbo, Guigo, & Breschi, 2018) was used to generate a sashimi plot of average reads across the *Myc* gene.

### qRT-PCR

RNA was isolated from equivalent numbers of wing imaginal discs (10 pairs for each genotype) using the Promega ReliaPrep RNA Tissue miniprep system and eluted in 20 µL nuclease-free water. RNA purity and integrity were assessed using an automated electrophoresis system (2200 TapeStation, Agilent Technologies). 5 µL of RNA was used for each cDNA synthesis (GoScript™ Reverse Transcription System kit, Promega). qPCR was performed using Fast SYBR Green Master Mix (Applied Biosystems) using the StepOnePlus Real-Time PCR System and Sequence Detection Systems in 96-well plates (Applied Biosystems, 95°C for 2 min, 40 cycles 95°C 1 s and 60°C 20 s). Amplicon specificity was verified by melt curve analysis. Average Ct values for two technical replicates were calculated for each sample. Multiple internal control genes were analyzed for stability and target gene expression was normalized to the mean of *cyp1* and *tubulin* or *cyp1* alone, selected for having high expression and little sample-to-sample variability as determined by RefFinder. Fold change was determined using the 2-ΔΔCT method.

Primers used:

Akap200 5’ GGCTACAAATGGCGAGGCTG 3’

5’ TTTCTCCGTTGGCCTGTTTCT 3’

chic 5’ TTTACCTTTCCGGCACAGACC 3’

5’ TGGAAACGATCACGGCTTGT 3’

dally 5’CATCATCACACCAGCAGCCT 3’

5’ GCCAATTCCAGGACGTGACT 3’

dlp 5’ TTTCCAAGCGAGAGGAATCG 3’

5’ ACCGAAGGGGACTCGCAATA 3’

emp 5’ GGACCCTACGTTTACAGCGA 3’

5’ TGTAGCTCAGCGTGCCATTG 3’

Exn 5’ CTTAAGGACCAAGCCGGCAA 3’

5’ AAGACAACACCAGCTCGACG 3’

fz2 5’ CGACTGCATGTGACACCAAAG 3’

5’ GGGCAATGTCGCCCATGAAA 3’

tok 5’ CGACTGCATGTGACACCAAAG 3’

5’ GGGCAATGTCGCCCATGAAA 3’

tubulin 5’ TCAGACCTCGAAATCGTAGC 3’

5’ AGCCTGACCAACATGGATAGAG 3’

cyp1 5’ TCGGCAGCGGCATTTCAGAT 3’

5’ TGCACGCTGACGAAGCTAGG 3’

### Adult wing analysis

Adult wings were mounted in paraffin oil. Adult wing size was determined for male wings that were imaged with an Olympus SZ51 binocular microscope, at 4x magnification using an Olympus DP20 camera. Wing size was measured by pixel count for the area posterior to wing vein L5, using Photoshop software CS5.

### Statistics

All statistical tests that were not part of the RNAseq or DamID analysis were performed with Graphpad Prism 7 using unpaired 2-tailed t-test with 95% confidence interval. In all figures, the error bars represent SD and significance represented according to the Graphpad classification * (P = 0.01–0.05), ** (P = 0.001–0.01), *** (P = 0.0001–0.001) and **** (P< 0.0001).

## Acknowledgments

The authors acknowledge the facilities and the scientific and technical assistance of the Biomolecular Resource Facility funded by Australian Cancer Research Foundation (ACRF), located in the John Curtin School of Medical Research. This work was supported by the National Health and Medical Research Council [APP1143008 to L.M.Q.] and National Institutes of Health [NIH GM25232 to J.T.L]. The content is solely the responsibility of the authors and does not necessarily represent the official views of the National Institutes of Health.

## Data Availability

DamID and RNAseq data are available via the Gene Expression Omnibus (GEO) using accession number GSE149721

---

**Supplemental Figure 1.**
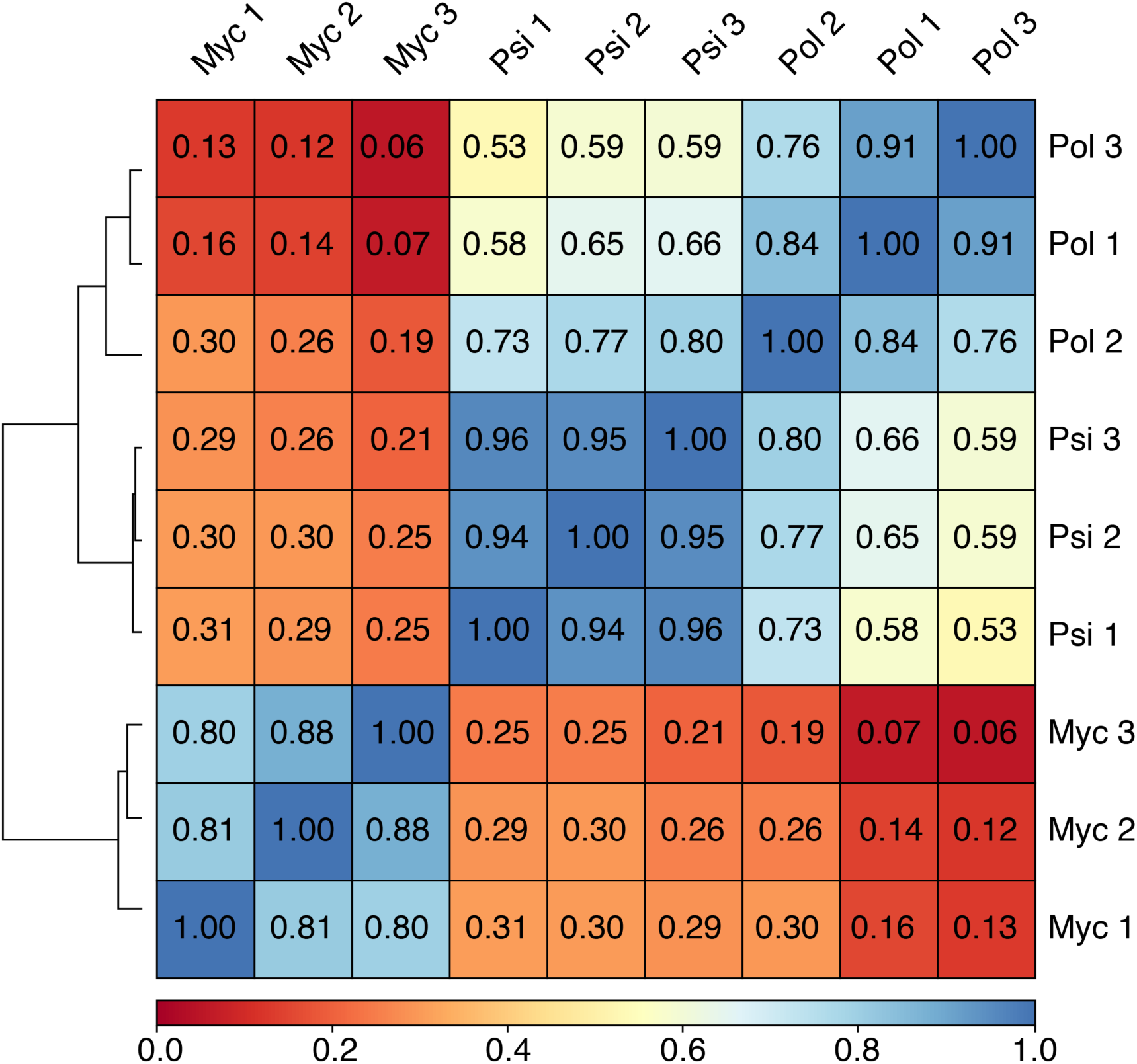
DamID sample correlation. Clustered heatmap showing values for Spearman correlation of individual Psi, Myc and Pol DamID samples to verify data quality and concordance between replicate samples.

**Supplemental Figure 2.**
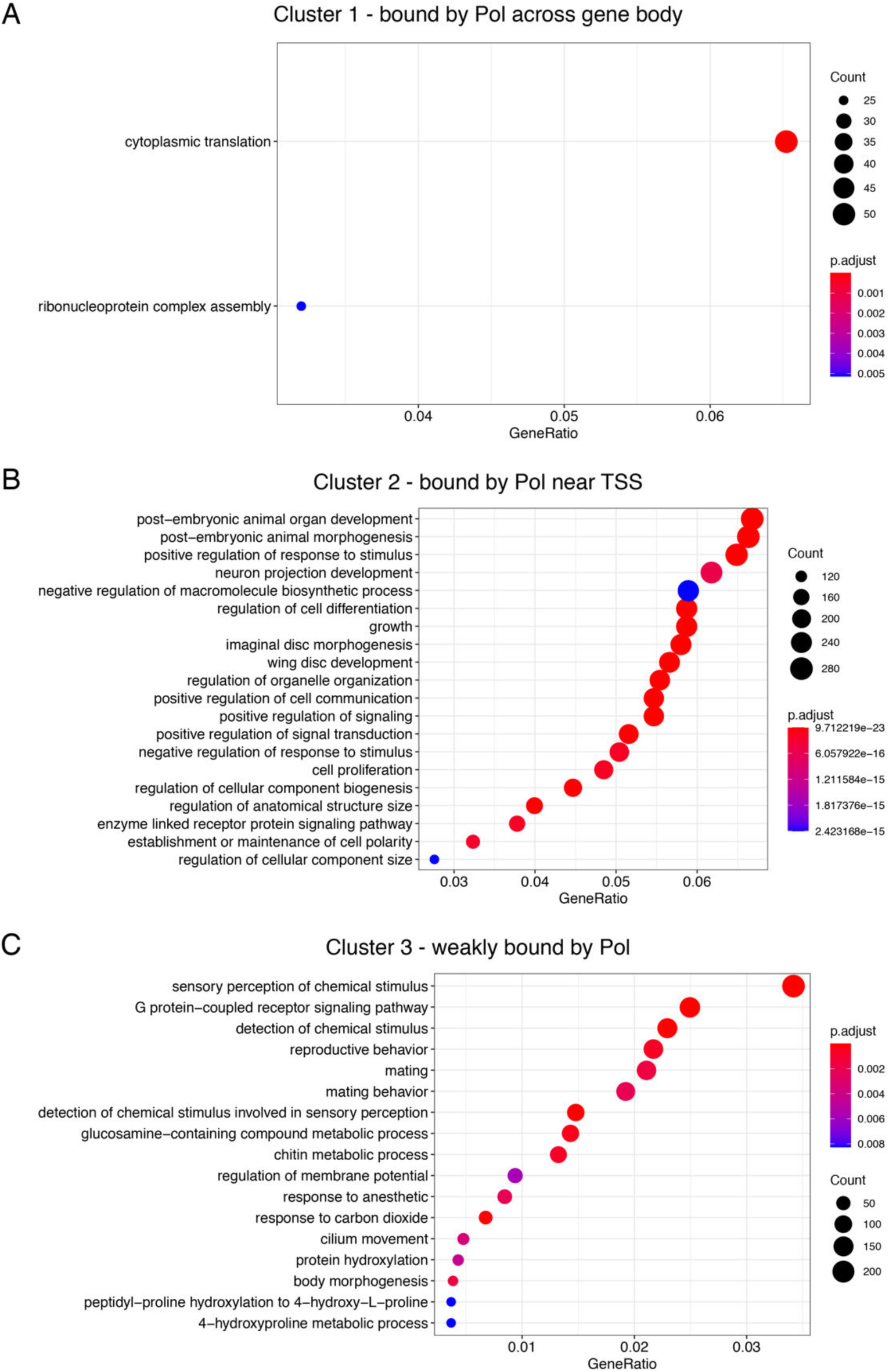
Ontology of genes showing similar transcriptional activity across 3 major clusters. Analysis of gene clusters shown in Figure 2A of the main text.

**Supplemental Figure 3.**
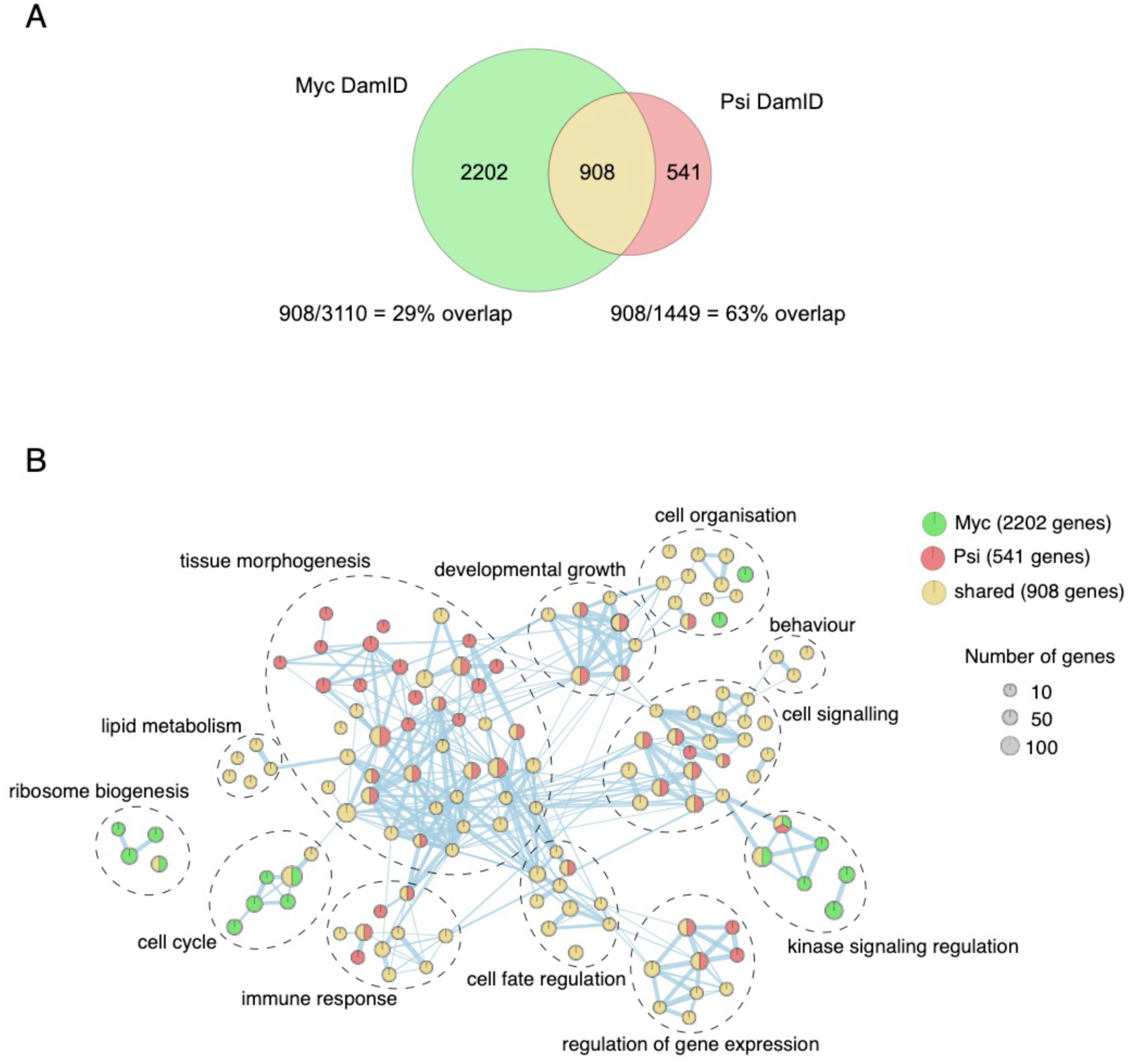
Ontology of genes with significant Psi and Myc binding peaks. (A) Intersection of Psi and Myc DamID target genes (FDR<0.01). (B) Network of gene ontologies enriched among genes bound by Psi alone (red), Myc alone (green) and both Psi/Myc (yellow).

**Supplemental Figure 4.**
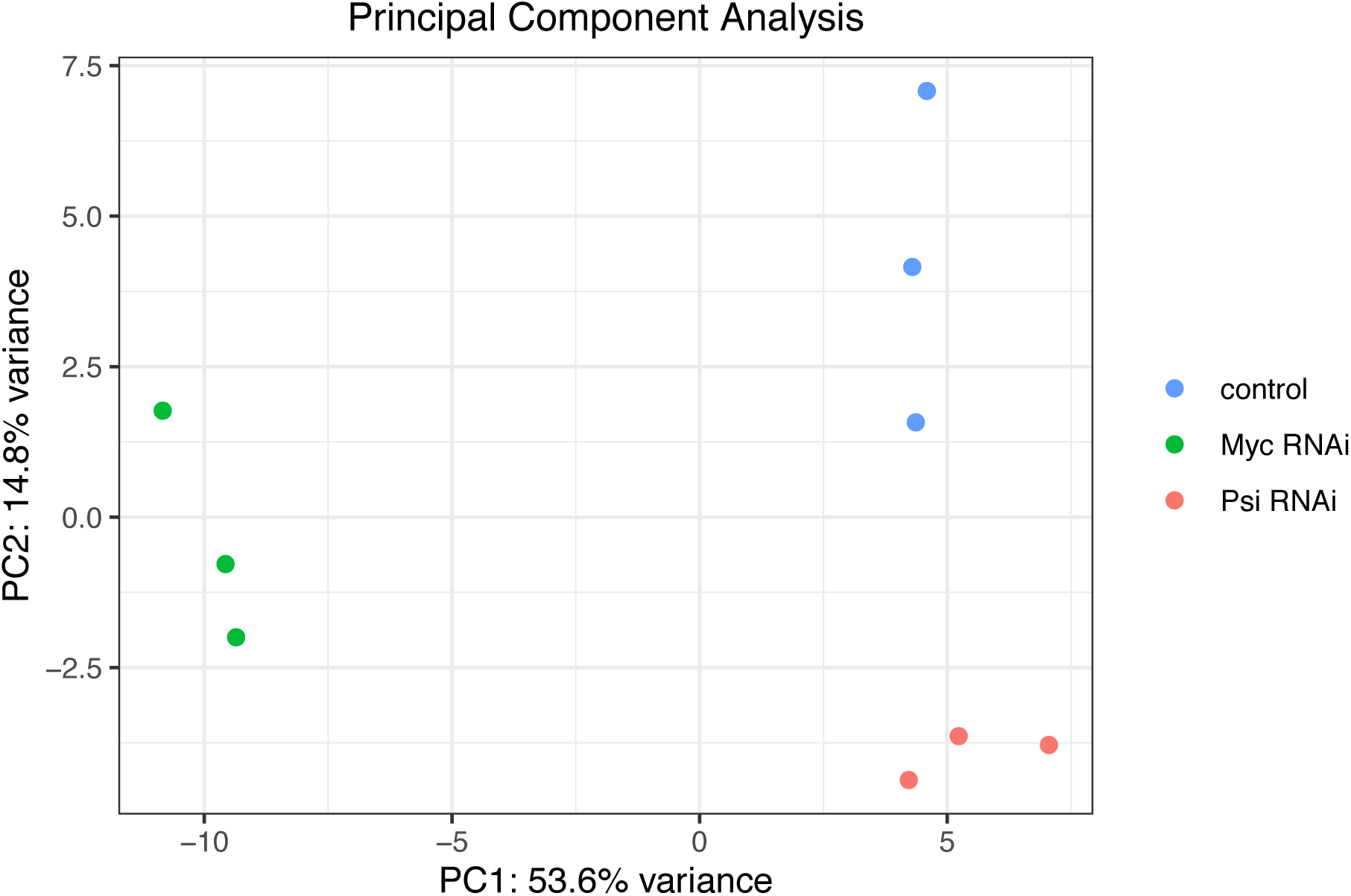
RNA-seq sample correlation. Principal component analysis of RNA-seq samples following expression of Psi RNAi or Myc RNAi in larval wing discs using tsGAL80; tub-GAL4, compared with control.

**Supplemental Figure 5.**
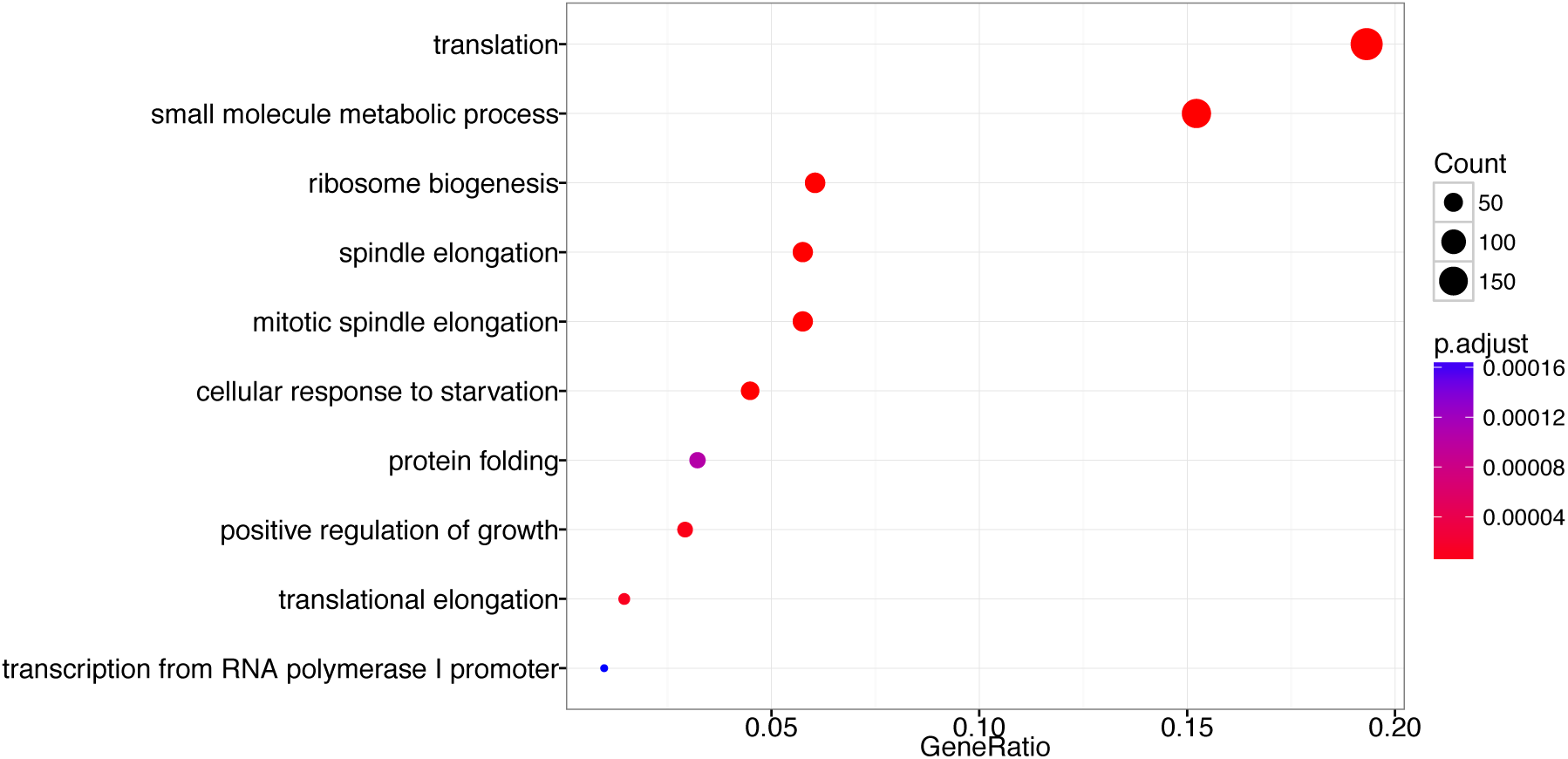
Genes downregulated after Myc depletion function in growth. Ontology analysis of genes downregulated after knockdown of Myc.

**Supplemental Figure 6.**
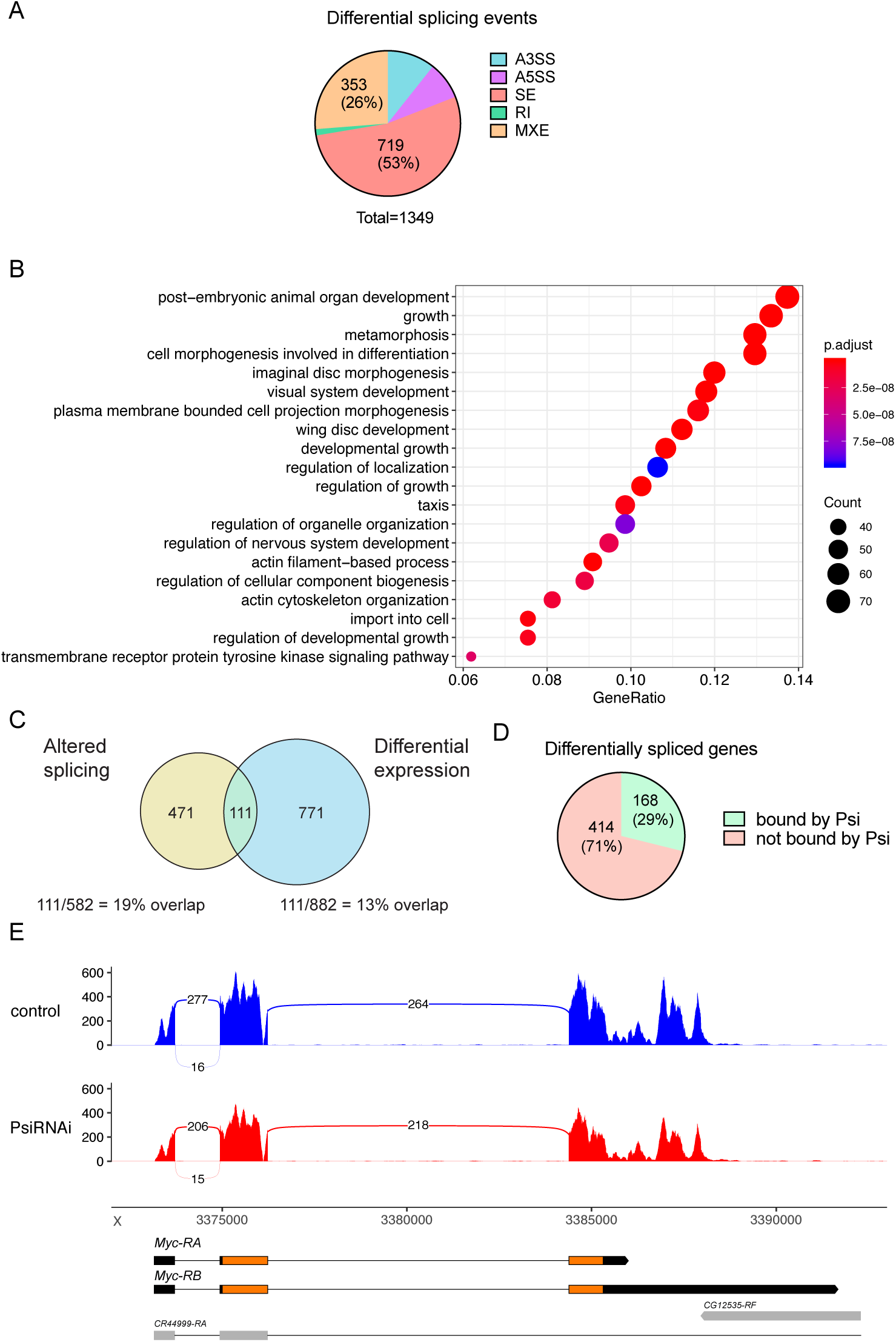
Splicing analysis of *Psi* RNAi RNA-seq. (A) Proportion of differential splicing events detected by rMATS at FDR<0.01: alternative 3’splice site (A3’SS), alternative 5’ splice site (A5’SS), skipped exon (SE), retained intron (RI) mutually exclusive exons (MXE). (B) Gene ontology analysis of genes with differential splicing detected by rMATS. (C) Intersection of differentially expressed genes with genes exhibiting altered splicing. (D) Proportion of differentially spliced genes that are bound by Psi. (E) Sashimi plot of the *Myc* gene shown as the average of three RNA-seq replicates.

**Supplemental Figure 7.**
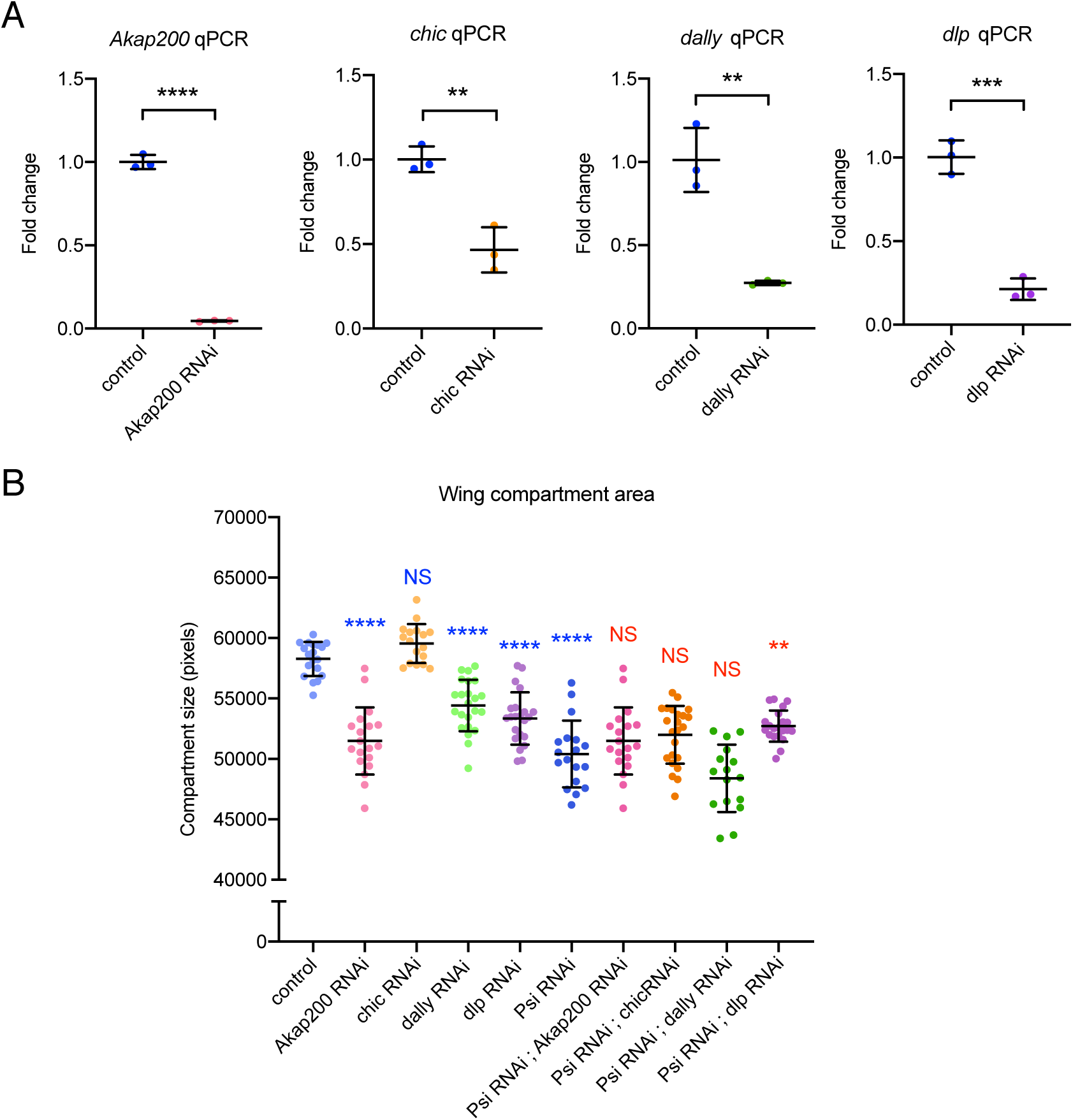
Targets negatively regulated by Psi not required for Psi-dependent growth. (A) qPCR of third instar larval wing discs 2 days after induction of RNAi transgenes (using tsGAL80; *tub*-GAL4) for Psi targets as labelled. (B) Quantification of the posterior compartment of the adult wing defined by the L5 vein. P-values were corrected for multiple testing using the Bonferroni method (blue stars indicate comparison to control; red stars indicate comparison to *Psi* RNAi). **** indicates p_adj_<0.0001, ** p_adj_=0.0024, NS=not significant.

## Supplemental Information

RpII18_gBlock sequence (lower case = Gibson assembly overlaps with *pTaDaG*):

ctcatctctgaagaggatctggccggcgcaCCGGCCGCCAAGCGCGTGAAGCTGGATGGCGCCGGCATGGATGATGCGGACTACGACAACGACGACGTTGGCGGCGATGACTTCGACGACGTCGACGAGGACGTGGACGAGGACATTAACCAGGAGGAGGAGGCGGACAACATCGAGATCATAGCTCCCGGTGGTGCGGGCGGAGGCGGTGTGCCCAAGTCCAAGCGCATTACCACAAAGTACATGACGAAATACGAGCGCGCCAGAGTTCTGGGCACACGAGCGCTTCAGATCGCCATGTGCGCACCCATCATGGTGGAGCTGGACGGGGAAACGGACCCCCTGCAGATCGCCATGAAAGAGCTGAAACAAAAGAAAATTCCCATCATCATCCGCCGATACCTGCCGGATCACTCCTACGAGGACTGGAGCATCGACGAGCTCATCATGGTGGACAACTAGgggtacctctagaggatctttgtgaaggaa

*Psi* ORF primers:

Fwd: CTCATCTCTGAAGAGGATCTGGCCGGCGCAATGAGCGACTTCCAGCAAC

Rev: TTCCTTCACAAAGATCCTCTAGAGGTACCCTCAGTGATTGTCGTTTTTGTGC

*Myc* ORF primers:

Fwd: CTCATCTCTGAAGAGGATCTGGCCGGCGCAATGGCCCTTTACCGCTCTG

Rev: TTCCTTCACAAAGATCCTCTAGAGGTACCCCTATCCACTAACCGAGCGC

